# Pan-Cancer Study Detects Novel Genetic Risk Variants and Shared Genetic Basis in Two Large Cohorts

**DOI:** 10.1101/635367

**Authors:** Sara R. Rashkin, Rebecca E. Graff, Linda Kachuri, Khanh K. Thai, Stacey E. Alexeeff, Maruta A. Blatchins, Taylor B. Cavazos, Douglas A. Corley, Nima C. Emami, Joshua D. Hoffman, Eric Jorgenson, Lawrence H. Kushi, Travis J. Meyers, Stephen K. Van Den Eeden, Elad Ziv, Laurel A. Habel, Thomas J. Hoffmann, Lori C. Sakoda, John S. Witte

**Affiliations:** Department of Epidemiology and Biostatistics, University of California, San Francisco, San Francisco, CA, USA; Division of Research, Kaiser Permanente Northern California, Oakland, CA, USA; Program in Biological and Medical Informatics, University of California, San Francisco, San Francisco, CA, USA; Department of Urology, University of California, San Francisco, San Francisco, CA, USA; Institute for Human Genetics, University of California, San Francisco, San Francisco, CA, USA; Department of Medicine, University of California, San Francisco, San Francisco, CA, USA; Helen Diller Family Comprehensive Cancer Center, University of California, San Francisco, San Francisco, CA, USA

**Author notes:** These authors contributed equally to this work (Sara R. Rashkin, Rebecca E. Graff). Co-senior / corresponding authors (Lori C. Sakoda, John S. Witte). Corresponding Authors: John S. Witte, Department of Epidemiology and Biostatistics, University of California, San Francisco, 1450 3rd Street, Room 388, San Francisco, CA 94158; Lori C. Sakoda, Division of Research, Kaiser Permanente Northern California, 2000 Broadway, Oakland, CA 94612.

## Abstract

Deciphering the shared genetic basis of distinct cancers has the potential to elucidate carcinogenic mechanisms and inform broadly applicable risk assessment efforts. However, no studies have investigated pan-cancer pleiotropy within single, well-defined populations unselected for phenotype. We undertook novel genome-wide association studies (GWAS) and comprehensive evaluations of heritability and pleiotropy across 18 cancer types in two large, population-based cohorts: the UK Biobank (413,870 European ancestry individuals; 48,961 cancer cases) and the Kaiser Permanente Genetic Epidemiology Research on Adult Health and Aging cohorts (66,526 European ancestry individuals; 16,001 cancer cases). The GWAS detected 21 novel genome-wide significant risk variants. In addition, numerous cancer sites exhibited clear heritability. Investigations of pleiotropy identified 12 cancer pairs exhibiting either positive or negative genetic correlations and 43 pleiotropic loci. We identified 158 pleiotropic variants, many of which were enriched for regulatory elements and influenced cross-tissue gene expression. Our findings demonstrate widespread pleiotropy and offer further insight into the complex genetic architecture of cross-cancer susceptibility.

## Introduction

The global burden of cancer is substantial, with an estimated 18.1 million individuals diagnosed each year and approximately 9.6 million deaths attributed to the disease.^1^ Efforts toward cancer prevention, screening, and treatment are thus imperative, but they require a more comprehensive understanding of the underpinnings of carcinogenesis than we currently possess. While studies of twins,^2^ families,^3^ and unrelated populations^4–6^ have demonstrated substantial heritability and familial clustering for many cancers, the extent to which genetic variation is unique versus shared across different types of cancer remains unclear.

Genome-wide association studies (GWAS) of individual cancers have identified loci associated with multiple cancer types, including 1q32 (*MDM4*)^7, 8^; 2q33 (*CASP8*-*ALS2CR12*)^9, 10^; 3q28 (*TP63*)^11, 12^; 4q24 (*TET2*)^13, 14^; 5p15 (*TERT-CLPTM1L*)^9, 12^; 6p21 (HLA complex)^15, 16^; 7p15^17^; 8q24^12, 18^; 11q13^18, 19^; 17q12 (*HNF1B*)^18, 20^; and 19q13 (*MERIT40*)^21^. In addition, recent studies have tested single nucleotide polymorphisms (SNPs) previously associated with one cancer to discover pleiotropic associations with other cancer types.^22–25^ Consortia, such as the Genetic Associations and Mechanisms in Oncology, have looked for variants and pathways shared by breast, colorectal, lung, ovarian, and prostate cancers.^26–30^ Comparable studies for other cancers— including those that are less common—have yet to be conducted.

In addition to individual variants, recent studies have evaluated genome-wide genetic correlations between pairs of cancer types.^4–6^ One evaluated 13 cancer types and found shared heritability between kidney and testicular cancers, diffuse large B-cell lymphoma (DLBCL) and osteosarcoma, DLBCL and chronic lymphocytic leukemia (CLL), and bladder and lung cancers.^4^ Another study of six cancer types found correlations between colorectal cancer and both lung and pancreatic cancers.^5^ In an updated analysis with increased sample size, the same group identified correlations of breast cancer with colorectal, lung, and ovarian cancers and of lung cancer with colorectal and head/neck cancers.^6^ While these studies provide compelling evidence for shared heritability across cancers, they lack data on several cancer types (e.g., cervix, melanoma, and thyroid).

Here, we present analyses of genome-wide SNP data with respect to 18 cancer types, based on 475,312 individuals of European ancestry from two large, independent, and contemporary cohorts unselected for phenotype – the UK Biobank (UKB) and the Kaiser Permanente Genetic Epidemiology Research on Adult Health and Aging (GERA) cohorts. We sought to detect novel risk SNPs and pleiotropic loci and variants and to estimate the heritability of and genetic correlations between cancer types. We then conducted *in-silico* functional analyses of pleiotropic variants to catalog biological mechanisms potentially shared across cancers. Leveraging the wealth of individual-level genetic and phenotypic data from both cohorts allowed us to extensively interrogate the shared genetic basis of susceptibility to different cancer types, with the ultimate goal of better understanding common genetic mechanisms of carcinogenesis and improving risk assessment.

## Results

### Genome-wide Association Analyses of Individual Cancers

We found 21 novel, independent variants that attained genome-wide significance at *P*<5×10^−8^ upon meta-analysis of the UKB and GERA results (**Table 1**) and an additional 9 genome-wide significant variants that were only genotyped or imputed in one cohort (**Supplementary Table 1**). In addition, we detected 308 independent signals with *P*<1×10^−6^ that confirmed risk SNPs identified by previous GWAS with *P*<5×10^−8^ (**Supplementary Table 2**). Of the 21 novel genome-wide significant associations from the meta-analyses, 9 were in known susceptibility regions for the cancer of interest but independent of previously reported variants. The remaining 12 were in regions not previously associated with the cancer of interest in individuals of European ancestry. Fourteen of the 21 novel SNPs exhibited pleiotropy in that they were in regions previously associated with at least one of the other cancer types evaluated in this study.

**Table 1.** Novel genome-wide significant loci from meta-analysis of UK Biobank (UKB) and Genetic Epidemiology Research on Adult Health and Aging (GERA) SNPs for each cancer site.

| Cancer Site | SNP | Chromosome | Position | Gene | REF/ALT* | MAF** |  | Odds Ratio (OR) |  |  |  |
| --- | --- | --- | --- | --- | --- | --- | --- | --- | --- | --- | --- |
|  |  |  |  |  |  | UKB | GERA | UKB | GERA | Meta | Meta P |
| Bladder | rs76088467<sup>†,+</sup> | 6 | 21795787 | *CASC15* | **G/A** | 0.025 | 0.030 | 0.67 | 0.59 | 0.64 | 2.34x10<sup>-8</sup> |
| Breast | rs6752414<sup>†,+</sup> | 2 | 121425339 | Intergenic | **T/C** | 0.077 | 0.083 | 0.89 | 0.87 | 0.88 | 1.81x10<sup>-9</sup> |
| Breast | rs8027730<sup>§,+</sup> | 15 | 49872585 | *FAM227B* | **A/C** | 0.48 | 0.48 | 1.06 | 1.08 | 1.06 | 2.68x10<sup>-8</sup> |
| Cervix | rs10175462<sup>§,+</sup> | 2 | 113988492 | *PAX8* | **A/G** | 0.36 | 0.37 | 1.16 | 1.08 | 1.15 | 7.71x10<sup>-14</sup> |
| Cervix | rs2856437<sup>†,+</sup> | 6 | 32157364 | *PBX2* | **A/G** | 0.063 | 0.047 | 0.76 | 0.88 | 0.77 | 1.24x10<sup>-15</sup> |
| Colon | rs71518872<sup>§,+</sup> | 8 | 103561978 | Upstream of *ODF1* | **G/C** | 0.015 | 0.017 | 0.65 | 0.61 | 0.64 | 1.27x10<sup>-8</sup> |
| Colon | rs8114643<sup>§</sup> | 20 | 7833046 | Intergenic | **G/A** | 0.14 | 0.15 | 0.83 | 0.84 | 0.83 | 2.10x10<sup>-9</sup> |
| Esophagus/Stomach | rs75460256<sup>†,+</sup> | 2 | 106687838 | *C2orf40* | **G/A** | 0.024 | 0.022 | 0.52 | 0.67 | 0.53 | 1.04x10<sup>-8</sup> |
| Kidney | rs112248293<sup>§,+</sup> | 15 | 61500352 | *RORA* | **A/G** | 0.024 | 0.025 | 0.53 | 0.62 | 0.55 | 3.36x10<sup>-9</sup> |
| Lung | rs10863899<sup>§</sup> | 1 | 211666218 | 5'UTR of *RD3* | **G/A** | 0.42 | 0.42 | 1.23 | 1.09 | 1.18 | 1.91x10<sup>-8</sup> |
| Lung | rs146099759<sup>§</sup> | 5 | 12883592 | Intergenic | **A/G** | 0.024 | 0.028 | 0.69 | 0.57 | 0.64 | 3.50x10<sup>-8</sup> |
| Lung | rs12543486<sup>†,+</sup> | 8 | 13012376 | *DLC1* | **C/T** | 0.17 | 0.16 | 1.30 | 1.18 | 1.26 | 3.51x10<sup>-8</sup> |
| Lymphocytic Leukemia | rs114490818<sup>†</sup> | 3 | 126099101 | Intergenic | **A/G** | 0.022 | 0.011 | 0.48 | 0.53 | 0.48 | 2.86x10<sup>-8</sup> |
| Lymphocytic Leukemia | rs61965473<sup>§,+</sup> | 13 | 95571786 | Intergenic | **T/C** | 0.023 | 0.023 | 0.52 | 0.44 | 0.49 | 3.95x10<sup>-8</sup> |
| Lymphocytic Leukemia | rs78378222<sup>†,+</sup> | 17 | 7571752 | 3'UTR of *TP53* | **G/T** | 0.012 | 0.014 | 0.44 | 0.34 | 0.40 | 1.89x10<sup>-9</sup> |
| Melanoma | rs9818780<sup>§,+</sup> | 3 | 156492758 | Intergenic | **C/T** | 0.49 | 0.48 | 0.92 | 0.89 | 0.91 | 3.16x10<sup>-8</sup> |
| Melanoma | rs12186662<sup>§</sup> | 5 | 90356197 | *ADGRV1* | **G/A** | 0.32 | 0.36 | 0.90 | 0.89 | 0.90 | 1.09x10<sup>-8</sup> |
| Melanoma | rs55797833<sup>†,+</sup> | 9 | 21995044 | 5'UTR OF *CDKN2A* | **G/T** | 0.023 | 0.021 | 1.71 | 1.72 | 1.71 | 6.71x10<sup>-12</sup> |
| Melanoma | rs78378222<sup>†,+</sup> | 17 | 7571752 | 3'UTR of *TP53* | **G/T** | 0.012 | 0.014 | 0.70 | 0.63 | 0.67 | 1.18x10<sup>-8</sup> |
| Rectum | rs145503185<sup>§</sup> | 9 | 23455764 | Intergenic | **C/T** | 0.013 | 0.018 | 0.57 | 0.50 | 0.55 | 4.36x10<sup>-8</sup> |
| Thyroid | 2:173859846_TA_T<sup>§</sup> | 2 | 173859846 | *RAPGEF4* | **T/TA** | 0.25 | 0.26 | 1.45 | 1.15 | 1.36 | 3.49x10<sup>-8</sup> |

\*REF is reference allele and ALT allele is effect allele; bold allele is minor allele
\*\*MAF = minor allele frequency calculated in all controls
§ Indicates SNPs in loci not previously associated with the cancer of interest in European ancestry
† Indicates SNPs in known susceptibility loci for cancer of interest in European ancestry but independent of previously reported variants (LD $r^2 < 0.1$ in Europeans)
+ Indicates SNPs in loci previously associated with at least one of the other cancers evaluated in this study in European ancestry

In sensitivity analyses for the 21 novel SNPs, we did not detect any material differences in effect estimates across categories of age at diagnosis, Surveillance, Epidemiology, and End Results Program (SEER) grade, or SEER stage (Heterogeneity P>0.05, corrected for multiple testing). In genome-wide sensitivity analyses restricted to incident cases (i.e., excluding prevalent cases), our findings were essentially unchanged (Heterogeneity P>0.05, corrected for multiple testing; **Supplementary Figure 1**). Similarly, genome-wide sensitivity analysis results for esophageal and stomach cancers separately were materially comparable to those for the two phenotypes combined (Heterogeneity P>0.05, corrected for multiple testing; **Supplementary Figure 2**).

### Genome-Wide Heritability and Genetic Correlation

Array-based heritability estimates across cancers ranged from *h^2^*=0.04 (95% CI: 0.00-0.13) for oral cavity/pharyngeal cancer to *h^2^*=0.26 (95% CI: 0.15-0.38) for testicular cancer (**Table 2**). For some of the cancers, our array-based heritability estimates were comparable to twin- or family-based heritability estimates^2, 3^ but were more precise. Several were also similar to array-based heritability estimates from consortia comprised of multiple studies.^4–6^ For example, our estimate for testicular cancer closely matches the previous family-based estimate of heritability (*h^2^*=0.25; 95% CI: 0.15-0.37),^3^ as well as a previous estimate of array-based heritability (*h*^2^=0.30; 95% CI: 0.08-0.51).^4^ For lung cancer, our estimated heritability of *h^2^*=0.15 (95% CI: 0.10-0.20) approaches the twin-based estimate of *h^2^*=0.18 (95% CI: 0.00-0.42),^2^ exceeds the array-based estimate from a study using the same methodology (*h*^2^=0.08; 95% CI: 0.05-0.10),^6^ and is comparable to an earlier array-based estimate using individual-level data (*h^2^*=0.21; 95% CI: 0.14-0.27).^4^ For rectal (*h^2^*=0.11; 95% CI: 0.07-0.16) and bladder (*h^2^*=0.08; 95% CI: 0.04-0.12) cancers, our heritability estimates are close to those from twin/family studies – *h^2^*=0.14 (95%CI: 0.00-0.50)^2^ and *h^2^*=0.07 (95% CI: 0.02-0.11),^3^ respectively. One of our highest heritability estimates was observed for thyroid cancer (*h^2^*=0.21; 95% CI: 0.09-0.33), a cancer that has not been evaluated in other array-based studies.

**Table 2.** Heritability estimates (*h^2^*) and 95% confidence intervals (CIs) for each cancer based on the union set of UK Biobank (UKB) and Genetic Epidemiology Research on Adult Health and Aging (GERA) SNPs, compared with previous estimates from other array-based and twin/family-based studies.

| Cancer Site | Array-Based Heritability |  |  | Twin/Family-Based Heritability |
| --- | --- | --- | --- | --- |
|  | Current Study | Jiang *et al.*<sup>a</sup> | Sampson *et al.*<sup>c</sup> | Mucci *et al.*<sup>d</sup> |
| Bladder | 0.08 (0.04-0.12) |  | 0.12 (0.09-0.16) | 0.07 (0.02-0.11)<sup>e</sup> |
| Breast | 0.10 (0.08-0.13) | 0.14 (0.12-0.16) | 0.10 (0.00-0.20)** | 0.31 (0.11-0.51) |
| Cervix | 0.07 (0.02-0.12) |  |  | 0.13 (0.06-0.15)<sup>e,+</sup> |
| Colon | 0.07 (0.04-0.10) | 0.09 (0.07-0.11)* |  | 0.15 (0.00-0.45) |
| Endometrium | 0.13 (0.07-0.18) |  | 0.18 (0.09-0.27) | 0.27 (0.11-0.43) |
| Esophagus/Stomach | 0.14 (0.07-0.21) |  | 0.38 (0.17-0.59)*** | 0.22 (0.00-0.55)<sup>++</sup> |
| Kidney | 0.09 (0.04-0.15) |  | 0.15 (0.02-0.27) | 0.38 (0.21-0.55) |
| Lung | 0.15 (0.10-0.20) | 0.08 (0.05-0.10) | 0.21 (0.14-0.27) | 0.18 (0.00-0.42) |
| Lymphocytic Leukemia | 0.14 (0.05-0.23) |  | 0.22 (0.16-0.28)<sup>†</sup> | 0.09 (0.09-0.16)<sup>e,+++</sup> |
| Melanoma | 0.08 (0.04-0.11) |  |  | 0.58 (0.43-0.73) |
| Non-Hodgkin's Lymphoma | 0.13 (0.03-0.23) |  | 0.09 (0.04-0.15)<sup>††</sup> | 0.10 (0.08-0.10)<sup>e</sup> |
| Oral Cavity/Pharynx | 0.04 (0.00-0.13) | 0.10 (0.05-0.14) |  | 0.09 (0.00-0.60) |
| Ovary | 0.07 (0.01-0.13) | 0.03 (0.02-0.05) |  | 0.39 (0.23-0.55) |
| Pancreas | 0.06 (0.00-0.18) | 0.05 (0.00-0.10)<sup>b</sup> | 0.10 (0.04-0.16) |  |
| Prostate | 0.16 (0.13-0.20) | 0.18 (0.14-0.22) | 0.38 (0.24-0.51) | 0.57 (0.51-0.63) |
| Rectum | 0.11 (0.07-0.16) |  |  | 0.14 (0.00-0.50) |
| Testis | 0.26 (0.15-0.38) |  | 0.30 (0.08-0.51) | 0.25 (0.15-0.37)<sup>e</sup> |
| Thyroid | 0.21 (0.09-0.33) |  |  | 0.53 (0.52-0.53)<sup>e</sup> |

\* Colorectal
\*\* Estrogen receptor negative (ER-)
\*\*\* For esophageal in Asian population (stomach in Asian population: $h^2 = 0.25$ [0.00-0.52])
† For chronic lymphocytic leukemia
†† For diffuse large B-cell lymphoma
+ For in situ (invasive: $h^2 = 0.22$ [0.14-0.27])
++ Stomach
+++ Age >15 years
<sup>a</sup> Taken from Jiang, et al. (2019),<sup>6</sup> 95% CI calculated from provided standard error
<sup>b</sup> Taken from Lindström, et al. (2017)<sup>5</sup>
<sup>c</sup> Taken from Sampson, et al. (2015)<sup>4</sup>
<sup>d</sup> Taken from Mucci, et al. (2016),<sup>2</sup> except where not included in analysis or 95% CI range was > 0.60; remaining taken from Czene, et al. (2002),<sup>3</sup> as marked
<sup>e</sup> Taken from Czene, et al. (2002),<sup>3</sup> family-based not twin

Among pairs of cancers, only colon and rectal cancers (*r_g_*=0.85, *P*=5.33×10^−7^) were genetically correlated at a Bonferroni corrected significance threshold of *P*=0.05/153=3.27×10^−4^ (**Figure 1a-b; Supplementary Table 3**). However, at a nominal threshold of *P*=0.05, we observed suggestive relationships between 11 other pairs. Seven pairs showed positive correlations: esophageal/stomach cancer was correlated with Non-Hodgkin’s lymphoma (NHL; *r_g_*=0.40, *P*=0.0089), breast (*r_g_*=0.26, *P*=0.0069), lung (*r_g_*=0.44, *P*=0.0035), and rectal (*r_g_*=0.32, *P*=0.024) cancers; bladder and breast cancers (*r_g_*=0.22, *P*=0.017); melanoma and testicular cancer (*r_g_*=0.23, *P*=0.028); and prostate and thyroid cancers (*r_g_*=0.23, *P*=0.013). The remaining four pairs showed negative correlations: endometrial and testicular cancers (*r_g_*=-0.41, *P*=0.0064); esophageal/stomach cancer and melanoma (*r_g_*=-0.27, *P*=0.038); lung cancer and melanoma (*r_g_*=-0.28, *P*=0.0048); and NHL and prostate cancer (*r_g_*=-0.21, *P*=0.012).

**Figure 1.**
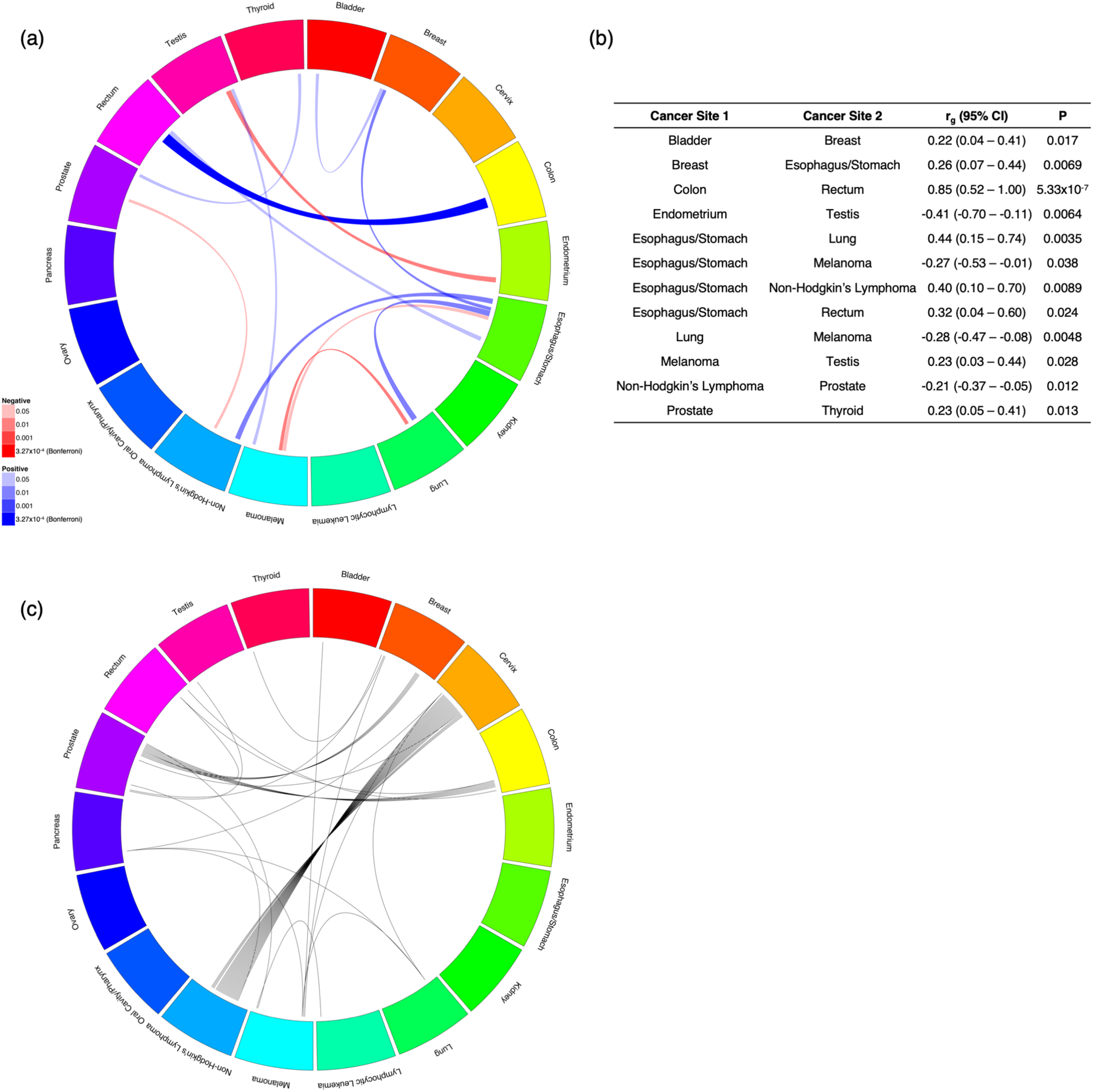
Cross-cancer genetic correlations (*r_g_*) calculated via LD-score regression (LDSC) and associated cancers from the locus-specific pleiotropy analysis. (a) Cancer pairs are connected if the genetic correlation had *P*<0.05, width of the line is proportional to magnitude of the point estimate, and shading is proportional to strength of association according to *P*, where the Bonferroni-corrected threshold is 0.05/153=3.27×10^−4^; (b) genetic correlation, 95% confidence interval (CI), and *P* for all cancer pairs with *P*<0.05; (c) cancer pairs are connected by a line (each line represents one region) if a region contains any SNPs associated with either cancer, where regions are formed around index SNPs with *P*<1×10^−6^ for any cancer and SNPs are added if they have *P*<1×10^−6^ for any cancer, are within 500kb of the index SNP, and have LD r^2^>0.5 with the index SNP.

### Locus-Specific Pleiotropy

We detected 43 pleiotropic regions associated with more than one cancer (*P*<1×10^−6^ for each cancer; regions defined using our linkage disequilibrium [LD] clumping procedure; see Methods; **Figure 1c; Supplementary Table 4**). Most were at known cancer pleiotropic loci: HLA (24 regions), 8q24 (10 regions), *TERT-CLPTM1L* (5 regions), and *TP53* (1 region). Twenty-two of the HLA regions were associated with both cervical cancer and NHL, and the remaining two were associated with (1) cervical and prostate cancers and (2) NHL and prostate cancer. Six regions in 8q24 were associated with prostate and colon cancers, and four were associated with prostate and breast cancers. Of the regions in *TERT-CLPTM1L*, two were associated with prostate cancer, one with breast cancer and one with testicular cancer; two were associated with melanoma, one with breast cancer and one with bladder cancer; and the last was associated with melanoma and cervical, lung, and pancreatic cancers. The *TP53* region, indexed by rs78378222, was associated with melanoma and lymphocytic leukemia. The remaining three pleiotropic regions were in loci previously associated with at least one cancer and were indexed by rs772695095 (*DIRC3* at 2q35; breast and thyroid cancers), rs11813268 (intergenic at 10q24.33; melanoma and prostate cancer), and rs6507874 (*SMAD7* at 18q21.1; colon and rectal cancers).

### Genome-wide Variant-Specific Pleiotropy

We assessed variant-specific pleiotropy by testing all variants genome-wide using the summary statistics for each cancer. We found 137 independent one-directional pleiotropic variants with at least two associated cancers, the same direction of effect for all associated cancers, and an overall pleiotropic *P*<1×10^−6^ (**Supplementary Table 5**), among which 85 attained genome-wide significance (*P*<5×10^−8^). Of the 137 one-directional pleiotropic variants, there were 45 for which the overall pleiotropic *P* was smaller than the *P* for each of the associated cancers, 17 of which attained genome-wide significance (**Figure 2**). While 134 of the 137 one-directional pleiotropic variants were in regions that have previously been associated with cancer, 113 were associated with at least one new cancer.

**Figure 2.**
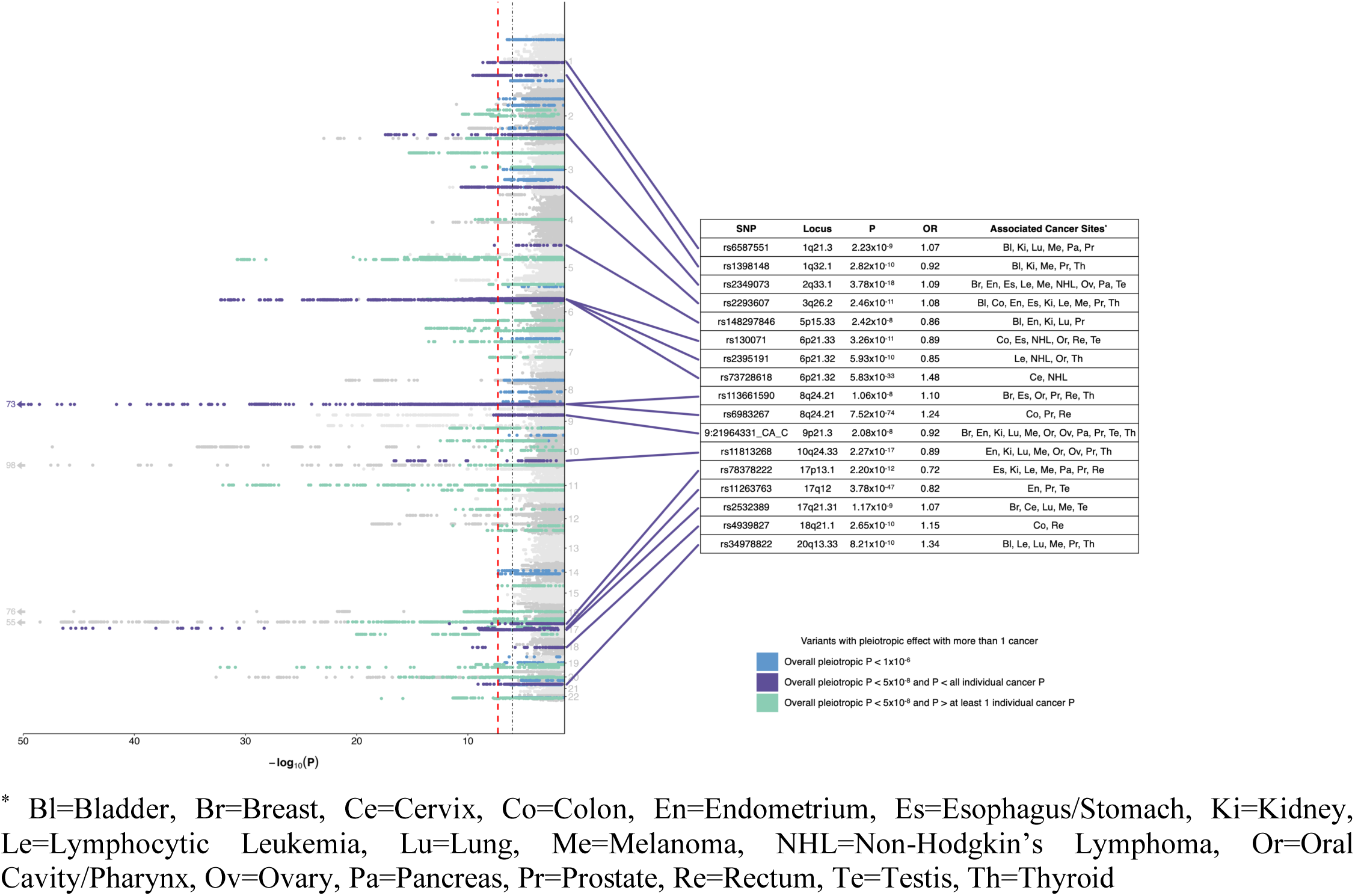
Manhattan plot displaying one-directional variant-specific pleiotropy, where the effect is maximized across all possible subsets of 18 cancers and assumes the same direction of effect across all selected cancers. The red dashed line represents the genome-wide significance threshold (*P* < 5×10^−8^), and the black dotted line represents a suggestive threshold (*P* < 1×10^−6^). Highlighted in purple are genome-wide significant loci where the overall pleiotropic *P* is less than all individual *P* for the selected cancers; details for the strongest signal at each locus are provided in the table, including the overall *P* and odds ratio (OR). Highlighted in green are the genome-wide significant loci where the overall pleiotropic *P* is greater than at least one of the individual *P* for the selected cancers, and highlighted in blue are loci with overall pleiotropic *P* < 1×10^−6^. All highlighted loci are independent of bidirectional SNPs with smaller overall *P*.

We also considered bidirectional pleiotropic associations, wherein the same allele for a given variant was associated with an increased risk for some cancers but a decreased risk for others. We found 21 such variants with *P*<1×10^−6^, all of which were independent from one another and from the one-directional pleiotropic variants (**Figure 3; Supplementary Table 6**). Fifteen attained genome-wide significance. There were 13 variants where the overall pleiotropic *P* was smaller than the *P* for the associated cancers, eight of which attained genome-wide significance. While 20 of the 21 bidirectional pleiotropic variants were in regions that have previously been associated with cancer, 10 were independent of known risk variants, and all 20 were associated with at least one new cancer. The SNP in a novel region (*SSPN* at 12q12.1) was rs10842692, which was associated with five cancers in one direction and nine cancers in the other.

**Figure 3.**
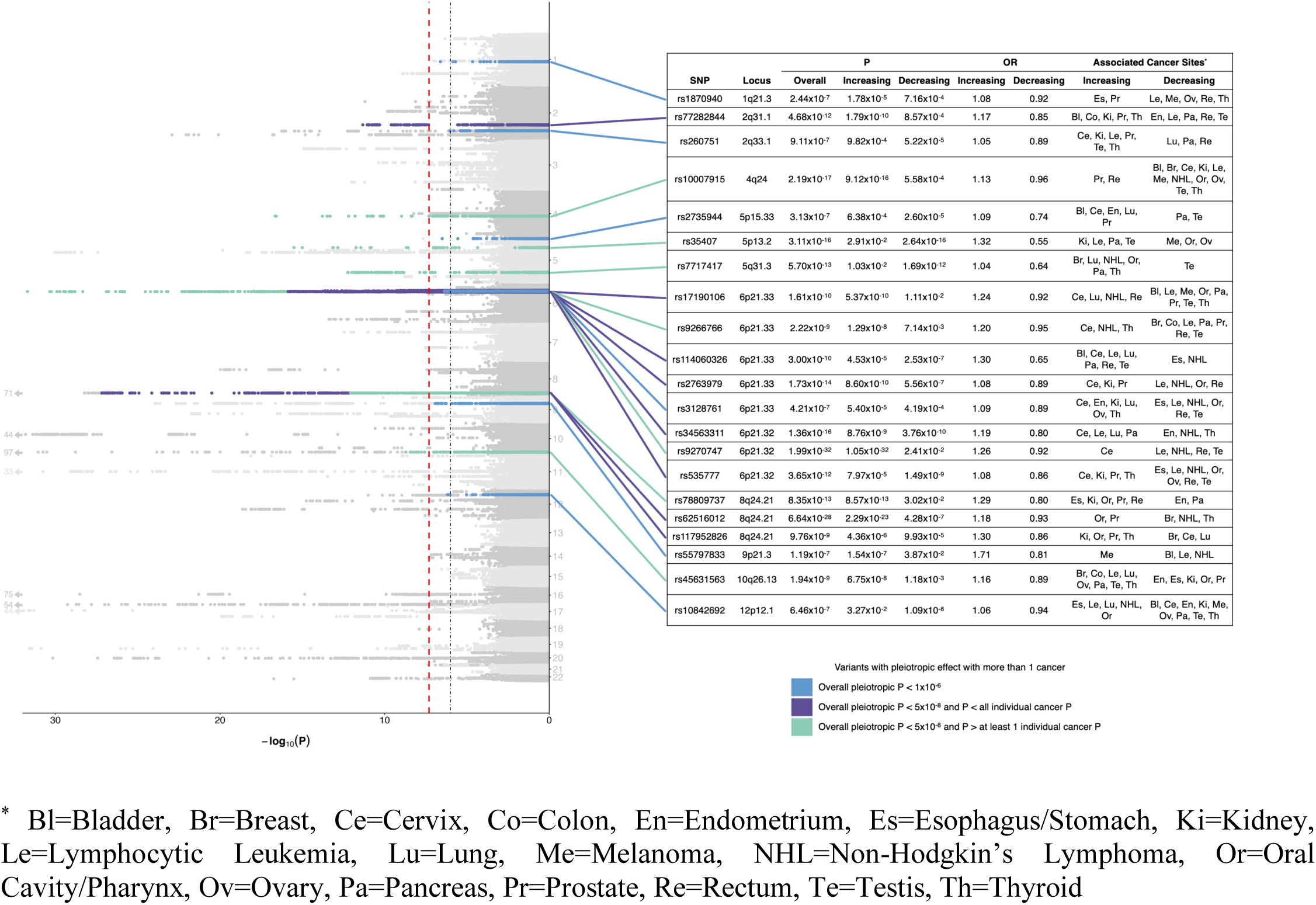
Manhattan plot displaying bidirectional variant-specific pleiotropy, where the effect is maximized across all possible subsets of 18 cancers and allows for different directions of effect for selected cancers. The red dashed line represents the genome-wide significance threshold (*P* < 5×10^−8^), and the black dotted line represents a suggestive threshold (*P* < 1×10^−6^). Highlighted are loci with overall pleiotropic *P* < 1×10^−6^, the two directional *P* < 0.05, and not in LD with a one-directional SNP with smaller *P*; loci in purple are genome-wide significant loci where the overall pleiotropic *P* is less than all individual *P* for the selected cancers, loci in green are genome-wide significant loci where the overall pleiotropic *P* is greater than at least one of the individual *P* for the selected cancers, and loci in blue have *P* < 1×10^−6^. Details for the strongest signal at each highlighted locus are provided in the table, including overall *P* and odds ratio (OR).

The number of one- and bidirectional SNPs shared by cancer pairs ranged from four (bladder and esophagus/stomach; colon and NHL) to 32 (pancreas and prostate; lymphocytic leukemia and prostate) (**Figure 4a; Supplementary Table 7**). For 19 cancer pairs, the shared associations had exclusively the same direction of effect (tabulating across both the one- and bidirectional analyses). For eight cancer pairs, most of the shared variants were associated in opposite directions.

**Figure 4.**
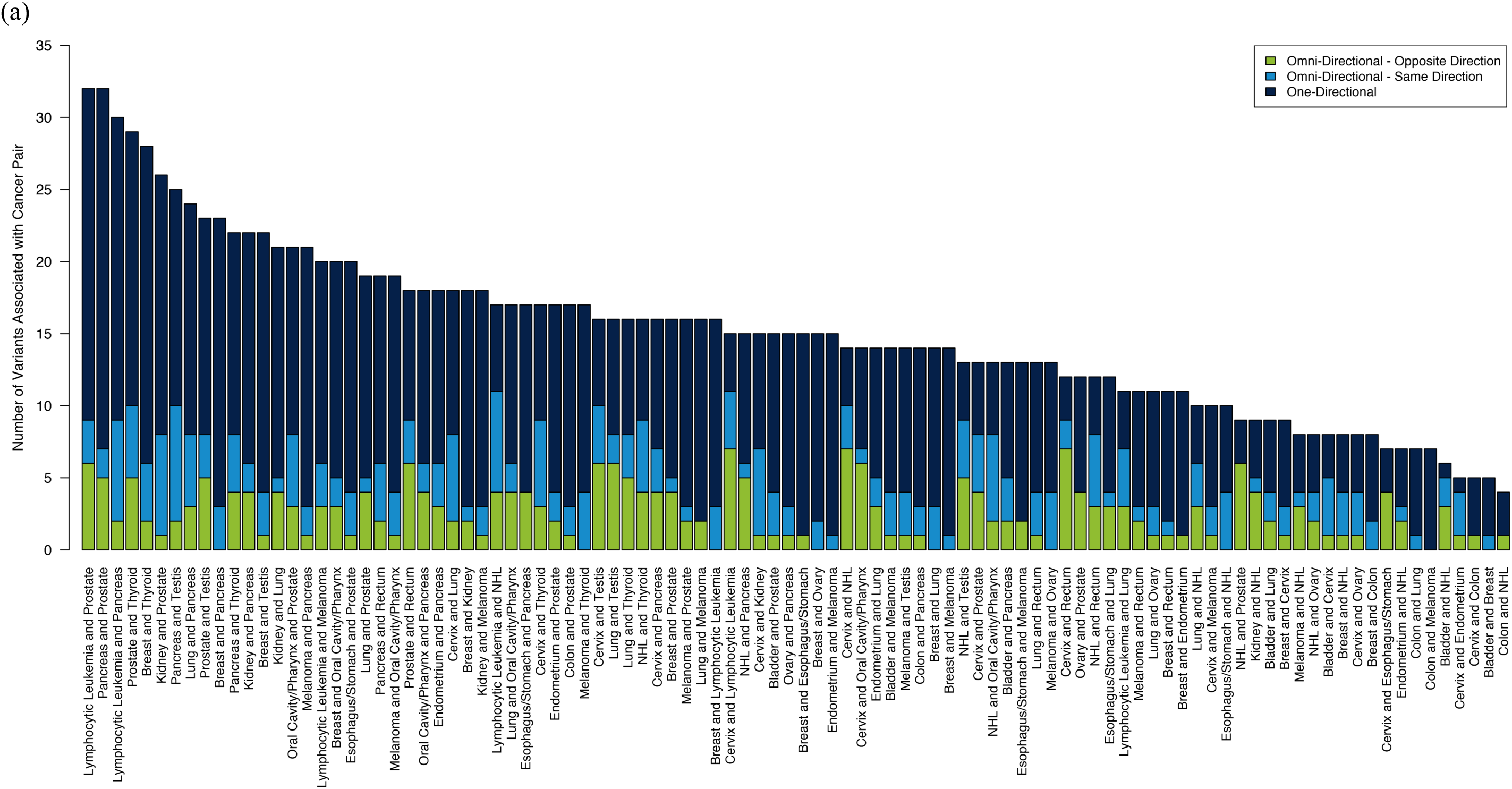
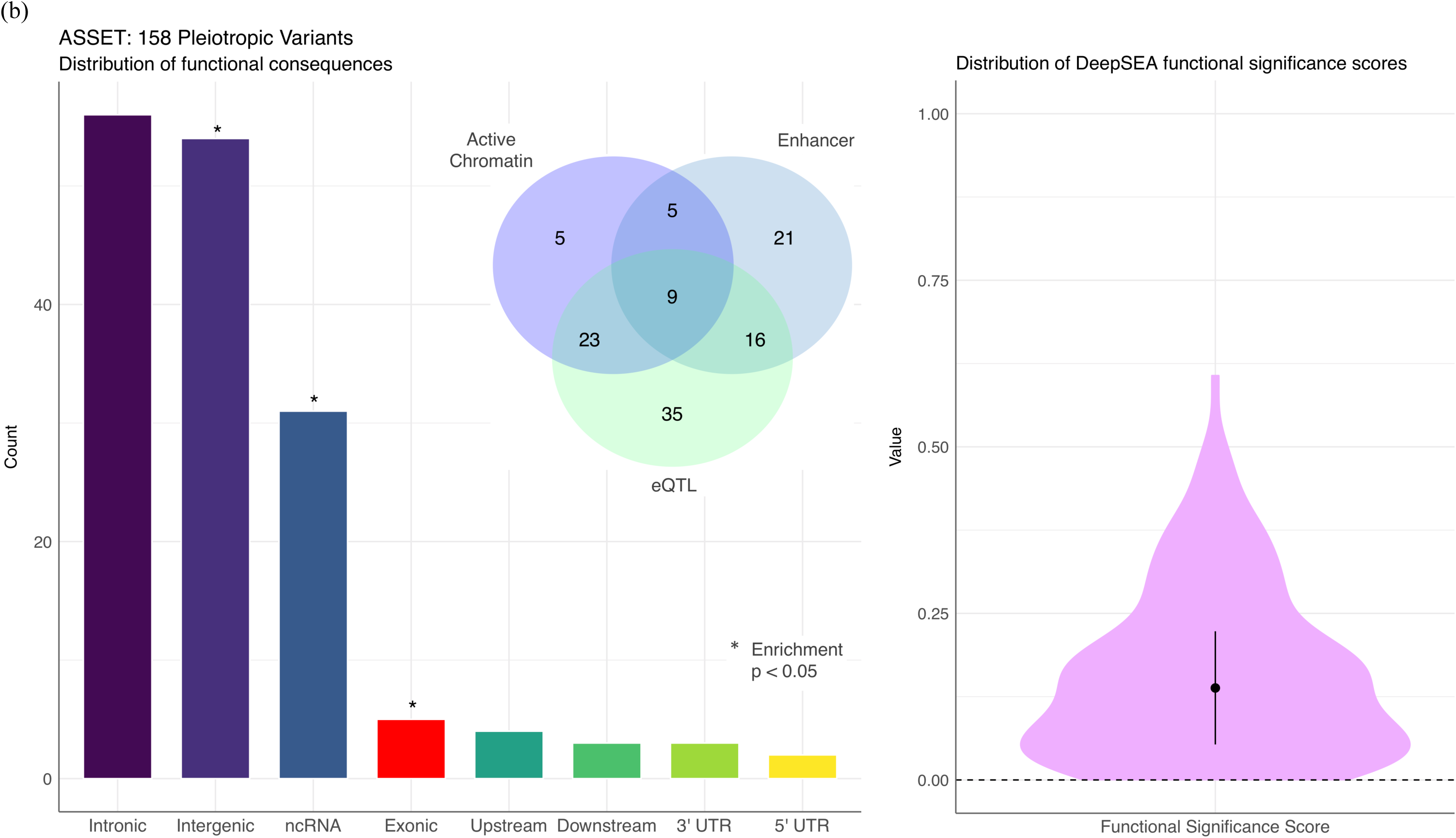
Summary of the cancer pairs associated with and the functional consequences of the 158 one- and bidirectional pleiotropic variants. (a) The number of pleiotropic variants (of the 158 one- and bidirectional variants with overall pleiotropic *P*<1×10^−6^) associated with each pair of cancers by type of pleiotropic effect for select cancer pairs. Variants were counted for a cancer pair if they were associated with both cancers using ASSET. For bidirectional variants, effects in the same direction and in opposite directions were tabulated separately. Included are all cancer pairs with any of the following cancer sites: breast, cervix, lung, melanoma, Non-Hodgkin’s Lymphoma (NHL), pancreas, and prostate. (b) The distribution of variant consequences and corresponding enrichment, calculated using Fisher’s exact test comparing the proportion of variants belonging to each functional class observed among the 158 ASSET variants to all variants in the UK Biobank. The Venn diagram summarizes the number of variants with specific regulatory elements, based on analyses of chromatin features from Roadmap and expression quantitative trait loci (eQTL) associations. DeepSEA functional significance score provides an integrated summary score based on evolutionary conservation and chromatin data, with 0 denoting variants most likely to be functional.

For each of the 158 SNPs showing either one- or bidirectional pleiotropy, we assessed whether the results differed according to age at diagnosis, SEER grade, or SEER stage for any of the associated cancers. After correcting for multiple testing, only one one-directional pleiotropic SNP showed heterogeneity across case subtypes. rs111362352-C was significantly positively associated with the risk of low grade prostate cancer in GERA, while it was not associated with high grade disease. These results are consistent with previous findings for this SNP (or SNPs in strong LD): the C allele has been associated with lower Gleason score, and it is located at *KLK3*, the prostate-specific antigen gene, which may reflect its previous association with lower grade screen-detected prostate cancer.^31, 32^

### Functional Characterization of Pleiotropic Variants

The biological significance of 158 pleiotropic variants was evaluated using *in-silico* annotation tools (**Supplementary Table 8**).^33–35^ Pleiotropic variants were enriched in intergenic and exonic regions, as well as non-coding RNA transcripts (*P*<0.018) (**Figure 4b)**. The distribution of DeepSea functional significance scores was skewed toward 0, indicating a higher likelihood of regulatory effects compared to a reference distribution of 1000 Genomes variants (**Figure 4b**). Suggestively functional variants (n=38, DeepSEA score<0.05) were also predicted to be pathogenic by Combined Annotation-Dependent Depletion (CADD; median score of 10.99, corresponding to the top 10% of deleterious substitutions). The top-ranked variant was rs3862792 in 11q13.3 (*CCND1*; synonymous; CADD=19.55), associated with prostate cancer and lymphocytic leukemia. Forty-two of the 158 pleiotropic variants were characterized by active chromatin states, 51 were classified as enhancers, and 83 had significant (FDR<0.05) effects on gene expression (**Figure 4b**). Nine variants belonged to all three classes (**Figure 4b)**, including rs10842692, the lead variant in one of our novel pleiotropic regions (*SSPN* at 12q12.1).

Consistent with hypothesized pleiotropy, 75.9% of the 83 expression quantitative trait loci (eQTLs) identified among the pleiotropic variants had more than one target tissue, and 71.1% influenced the expression of more than one gene (**Supplementary Figure 3**), for a total of 586 significant SNP-gene pairs. The most common expression tissues for eQTLs among pleiotropic variants were whole blood (63%), followed by subcutaneous adipose (9.9%), and esophageal (5.3%). Regulatory effects mediated by chromatin looping were observed for 46 variants, which clustered in the HLA region (10 variants), 8q24 (3 variants), and 11q13.3 (3 variants; **Supplementary Figure 4**). Three enhancer-promoter links were also identified in 6p21.23 (rs535777, rs73728618) and 22q13.2 (rs5759167, *PACSIN2* promoter; **Supplementary Figure 4**). Notably, rs5759167 is also an eQTL for *PACSIN2* in whole blood (*P*=9.89×10^−14^).

Genes represented by pleiotropic variants were significantly enriched for 67 pathways (**Supplementary Table 9**). Top-ranking pathways included inflammatory and immune response mechanisms (cytokines and chemokine targets of PSMD4: *p*=3.06×10^−8^; interferon-γ signaling in endothelial cells: *p*=1.16×10^−6^; antigen processing and presentation: *p*=2.53×10^−6^) and cell cycle checkpoint controls (G1/S: *p*=6.39×10^−6^). There was also enrichment for PML targets with promoters bound by Myc (*p*=3.27×10^−7^), transcription-factor networks implicated in neoplastic transformation, including FOXM1 (*p*=2.74×10^−5^), and p53 signaling (*p*=2.35×10^−4^).

## Discussion

In this study of cancer pleiotropy in two large cohorts, we found multiple lines of evidence for a shared genetic basis of several cancer types. By characterizing pleiotropy at the genome-wide, locus-specific, and variant-specific levels for a large number of cancer sites, we generated several novel insights into cancer susceptibility. Specifically, we detected 21 novel genome-wide significant risk variants across the 18 individual cancers. We also detected 158 variants displaying one- or bidirectional pleiotropy that were enriched for a number of regulatory functions that reflect hallmarks of carcinogenesis.

One notable finding from our cervical cancer GWAS was rs10175462 in *PAX8* on 2q13, which, to our knowledge, is the first genome-wide significant cervical cancer risk SNP identified outside of the HLA region in a European ancestry population.^15^ In a candidate SNP study of *PAX8* eQTLs in a Han Chinese population, two variants in LD with rs10175462 in Europeans (rs1110839, *r^2^*=0.33; rs4848320, *r^2^*=0.34) were suggestively associated with cervical cancer risk in the same direction.^36^ Several GWAS findings also provided evidence of pleiotropy, in that novel risk variants for one cancer had been previously associated with one or more other cancers. For instance, rs9818780 was associated with melanoma and has been implicated in sunburn risk.^37^ This intergenic variant is an eQTL for *LINC00886* and *METTL15P1* in skin tissue. The former gene has previously been linked to breast cancer,^38^ and both genes have been implicated in ovarian cancer.^39^ Beyond novel associations, our GWAS detected 308 independent associations with P<1×10^−6^ that confirmed signals identified in previous GWAS with P<5×10^−8^. This finding strengthened our confidence in using our genome-wide summary statistics for subsequent analyses of cancer pleiotropy.

In evaluating pairwise genetic correlations between the 18 cancer types, we observed the strongest signal for colon and rectal cancers – an expected relationship consistent with findings from a twin study.^40^ We also identified several novel cancer pairs for which the genetic correlations were nominally significant. One pair supported by previous evidence is melanoma and testicular cancer; some studies have found that individuals with a family history of the former are at an increased risk for the latter.^41, 42^ Esophageal/stomach cancer was a component of five correlated pairs – with melanoma, NHL, and breast, lung, and rectal cancers. Despite some similarities between esophageal and stomach cancers, testing them as a combined phenotype may have inflated the number of correlated cancers.

Our genetic correlation results contrast with previous consortia-based findings;^4–6^ we did not find several correlations that they did and found others that they did not. The differences may be partly due to a smaller number of cases in our cohorts for some sites. Further studies with larger sample sizes are necessary to validate our correlations, as those that did not attain Bonferroni-corrected significance may have been due to chance. However, we achieved comparable or higher cancer-specific heritability estimates for breast, colon, and lung cancers, which suggests that differences in study design may also play a role. Previous analyses aggregated case-control studies recruited during different time periods. While such meta-analyses can be effective at reducing residual population stratification, our extensive quality control processes also seemingly mitigated population stratification; the mean λ_GC_ across the 18 cancers was 1.02 (standard deviation=0.027). Moreover, our design allowed for the assessment of cross-cancer relationships in the same set of individuals, and to examine several cancers that have not been previously studied in large consortia.

The assessment of pleiotropy at the locus level confirmed previously reported associations at 5p15.33, HLA, and 8q24.^9, 12, 15, 16, 18^ Out of the 43 pleiotropic loci that we identified, over half, all in the HLA locus, were associated with cervical cancer and NHL. The two cancers were weakly negatively correlated in the two cohorts combined and nominally significantly negatively correlated in the UKB alone (**Supplementary Table 10**). The difference may reflect better coverage and imputation of the HLA region in the UKB than in GERA. Other findings support a pleiotropic role of several loci previously associated with specific cancers in separate studies. For example, we validated previous results showing that *DIRC3* is associated with breast and thyroid cancers.^14, 43^ Additionally, the intergenic region surrounding rs11813268 on 10q24.33 has not been previously associated with melanoma or prostate cancer (as it was in our study), although associations with other cancers have been reported, including kidney, lung, and thyroid cancers.^39, 44–47^ *SMAD7* has been previously linked to colorectal cancer,^48^ and we confirmed its association with colon and rectal cancers separately.

Variant-specific analyses provided further evidence in support of locus-specific cancer pleiotropy, including validation of previously reported signals at 1q32^7, 8^ and 2q33^9, 10^ (*ALS2CR12*). Interestingly, our lead 1q32 variant (rs1398148) maps to *PIK3C2B* and is in LD (*r*^2^>0.60) with known *MDM4* cancer risk variants,^7, 8^ suggesting that the 1q32 locus may be involved in modulating both p53-and PI3K-mediated oncogenic pathways. The 158 pleiotropic variants (with overall pleiotropic *P*<1×10^−6^) mapped to a total of 78 genomic locations, which included all of the regions identified from the locus-specific analyses. Although 154 of the 158 variants showing one- or bidirectional pleiotropic associations are in regions previously associated with cancer, 133 of the 154 were associated with at least one new cancer. One variant (rs10842692) associated with 14 cancers in the bidirectional analysis mapped to a novel region at 12p12.1 and is an active enhancer and an eQTL for *SSPN* in multiple tissue types, including adipose. *SSPN* has been linked to waist circumference,^49^ suggesting that increased adiposity may be one plausible mechanism underlying the pleiotropic associations observed for this locus.

Out of 158 total variants identified from the variant-specific pleiotropy analyses, 20 were in 8q24 and 19 were in the HLA region. Different distributions of one- and bidirectional results highlight patterns of directional pleiotropy: of the 19 HLA variants, eight were bidirectional, while only three of the 20 variants in 8q24 were bidirectional. The HLA region is critical for innate and adaptive immune response and has a complex relationship with cancer risk. Heterogeneous associations with HLA haplotypes have been reported for different subtypes of NHL^50^ and lung cancer,^51^ suggesting that relevant risk variants are likely to differ within, as well as between, cancers. Studies have further demonstrated that somatic mutation profiles are associated with HLA class I^52^ and class II alleles.^53^ Specifically, mutations that create neoantigens more likely to be recognized by specific HLA alleles are less likely to be present in tumors from patients carrying such alleles. It is thus possible that some of the positive and negative pleiotropy we identified is related to mutation type. These results reinforce the importance of the immune system playing a role in cancer susceptibility.

In contrast to the HLA region, the majority of the 8q24 pleiotropic variants had the same direction of effect for all associated cancers, implying the existence of shared genetic mechanisms driving tumorigenesis across sites. The proximity of the well-characterized *MYC* oncogene makes it a compelling candidate for such a consistent, one-directional effect. It could work via regulatory elements, such as acetylated and methylated histone marks.^54^ Consistent with this hypothesis, we observed heritability enrichment^55^ for variants with the H3K27ac annotation for breast (*P* = 3.09×10^−4^), colon (*P* = 4.44×10^−4^), prostate (*P* = 2.74×10^−5^), and rectal (*P* = 0.036) cancers – all of which share susceptibility variants in 8q24, according to our analyses and previous studies.^54^

*In-silico* analyses found the 158 pleiotropic variants to be enriched across multiple regulatory domains and highlighted functional features relevant to cancer pleiotropy. The 11q13.3 region includes rs3862792, which is predicted to be in the top 1% of deleterious substitutions in the genome.^35^ Although rs3862792 has been previously linked to prostate cancer,^56^ our results suggest it may also be relevant for lymphocytic leukemia. *CCND1* is a hallmark cancer oncogene that plays a role in cell cycle transitions, cell invasion, and cell migration, making rs3862792 a highly plausible candidate for cross-cancer effects. Three pleiotropic variants in 11q13.3 mapped to enhancer regions, including an eQTL for *TPCN2*, which is part of a signaling pathway controlling the angiogenic response to VEGF.^57^ The 22q13.2 region is indexed by rs5759167, an intergenic variant linked to prostate and lung cancers in our analysis. Its pleiotropic effects are likely mediated by regulation of *PACSIN2*, which codes for a cyclin D1 binding partner that serves as a brake for *CCND1*-mediated cellular migration.^58^ This is consistent with our observation that that the risk-increasing G-allele was associated with increased *PACSIN2* expression in whole blood.^59^ Lastly, our pathway analysis indicated that pleiotropic variants as a group are enriched for canonical signaling pathways that control cell-cycle progression, apoptosis, and immune-related functions, the dysregulation of which is a hallmark of cancer.

It is important to acknowledge some limitations of our study. First, counts for some of the cancer types were limited. However, small sample sizes are partially offset by the advantages of using two population-based cohorts. Second, due to the complexity of the LD structure in the HLA region, we may have overestimated the number of distinct, independent signals. Slight overestimation, however, does not affect our overall conclusions regarding the pleiotropic nature of this region. Third, our analyses included both prevalent and incident cases. Nevertheless, sensitivity analyses restricted to incident cancers yielded comparable results. Fourth, we grouped esophageal and stomach cancers despite some differences in their risk factor profiles. However, there is precedent for using a composite phenotype,^60^ and analyses of stomach and esophageal tumors suggest that they have many overlapping molecular features.^61, 62^ In addition, sensitivity analyses for each cancer alone gave similar results, suggesting that they may have similar genetic bases despite having different environmental risk factors. Finally, we focused solely on individuals of European ancestry. Further analyses are needed to accurately assess patterns of pleiotropy in non-Europeans.

The characterization of pleiotropy is fundamental to understanding the genetic architecture of cross-cancer susceptibility and its biological underpinnings. The availability of two large, independent cohorts provided an unprecedented opportunity to efficiently evaluate the shared genetic basis of many cancers, including some not previously studied together. The result was a multifaceted assessment of common genetic factors implicated in carcinogenesis, and our findings illustrate the importance of investigating different aspects of cancer pleiotropy. Broad analyses of genetic susceptibility and targeted analyses of specific loci and variants may both contribute insights into different dimensions of cancer pleiotropy. Future studies should consider the contribution of rare variants to cancer pleiotropy and aim to elucidate the functional pathways mediating associations observed at pleiotropic regions. Such research, combined with our findings, has the potential to inform drug development, risk assessment, and clinical practice toward reducing the burden of cancer.

## Methods

### Study Populations and Phenotyping

The UKB is a population-based cohort of 502,611 individuals in the United Kingdom. Study participants were aged 40 to 69 at recruitment between 2006 and 2010, at which time all participants provided detailed information about lifestyle and health-related factors and provided biological samples.^63^ GERA participants were drawn from adult Kaiser Permanente Northern California (KPNC) health plan members who provided a saliva sample for the Research Program on Genes, Environment and Health (RPGEH) between 2008 and 2011. Individuals included in this study were selected from the 102,979 RPGEH participants who were successfully genotyped as part of GERA and answered a baseline survey concerning lifestyle and medical history.^64, 65^

Cancer cases in the UKB were identified via linkage to various national cancer registries established in the early 1970s.^63^ Data in the cancer registries are compiled from hospitals, nursing homes, general practices, and death certificates, among other sources. The latest cancer diagnosis in our data from the UKB occurred in August 2015. GERA cancer cases were identified using the KPNC Cancer Registry, including all diagnoses captured through June 2016. Following SEER standards, the KPNC Cancer Registry contains data on all primary cancers (i.e., cancer diagnoses that are not secondary metastases of other cancer sites; excluding non-melanoma skin cancer) diagnosed or treated at any KPNC facility since 1988.

In both cohorts, individuals with at least one recorded prevalent or incident diagnosis of a borderline, in situ, or malignant primary cancer were defined as cases for our analyses. Individuals with multiple cancer diagnoses were classified as a case only for their first cancer. For the UKB, all diagnoses described by International Classification of Diseases (ICD)-9 or ICD-10 codes were converted into ICD-O-3 codes; the KPNC Cancer Registry already included ICD-O-3 codes. We then classified cancers according to organ site using the SEER site recode paradigm.^66^ We grouped all esophageal and stomach cancers and, separately, all oral cavity and pharyngeal cancers to ensure sufficient statistical power. The 18 most common cancer types (except non-melanoma skin cancer) were examined. Testicular cancer data were obtained from the UKB only due to the small number of cases in GERA.

Controls were restricted to individuals who had no record of any cancer in the relevant registries, who did not self-report a prior history of cancer (other than non-melanoma skin cancer), and, if deceased, who did not have cancer listed as a cause of death. For analyses of sex-specific cancer sites (breast, cervix, endometrium, ovary, prostate, and testis), controls were restricted to individuals of the appropriate sex.

### Quality Control

For the UKB population, genotyping was conducted using either the UKB Axiom array (436,839 total; 408,841 self-reported European) or the UK BiLEVE array (49,747 total; 49,746 self-reported European).^63^ The former is an updated version of the latter, such that the two arrays share over 95% of their marker content. UKB investigators undertook a rigorous quality control (QC) protocol.^63^ Genotype imputation was performed using the Haplotype Reference Consortium as the main reference panel and the merged UK10K and 1000 Genomes phase 3 reference panels for supplementary data.^63^ Ancestry principal components (PCs) were computed using *fastPCA*^67^ based on a set of 407,219 unrelated samples and 147,604 genetic markers.^63^

For GERA participants, genotyping was performed using an Affymetrix Axiom array (Affymetrix, Santa Clara, CA, USA) optimized for individuals of European race/ethnicity. Details about the array design, estimated genome-wide coverage, and QC procedures have been published previously.^65, 68^ The genotyping produced high quality data with average call rates of 99.7% and average SNP reproducibility of 99.9%. Variants that were not directly genotyped (or that were excluded by QC procedures) were imputed to generate genotypic probability estimates. After pre-phasing genotypes with SHAPE-IT v2.5,^69^ IMPUTE2 v2.3.1 was used to impute SNPs relative to the cosmopolitan reference panel from 1000 Genomes.^70–72^ Ancestry PCs were computed using Eigenstrat v4.2, as previously described.^64^

For both cohorts, analyses were limited to self-reported European ancestry individuals for whom self-reported and genetic sex matched. To further minimize potential population stratification, we excluded individuals for whom either of the first two ancestry PCs fell outside five standard deviations of the mean of the population. Based on a subset of genotyped autosomal variants with minor allele frequency (MAF) ≥0.01 and genotype call rate ≥97%, we excluded samples with call rates <97% and/or heterozygosity more than five standard deviations from the mean of the population. With the same subset of SNPs, we used KING^73^ to estimate relatedness among the samples. We excluded one individual from each pair of first-degree relatives, first prioritizing on maximizing the number of the cancer cases relevant to these analyses and then maximizing the total number of individuals in the analyses. Our study population ultimately included 413,870 UKB participants and 66,526 GERA participants. We excluded SNPs with imputation quality score <0.3, call rate <95% (alternate allele dosage required to be within 0.1 of the nearest hard call to be non-missing; UKB only), Hardy-Weinberg equilibrium *P* among controls <1×10^−5^, and/or MAF <0.01, leaving 8,876,519 variants for analysis for the UKB and 8,973,631 for GERA.

### Genome-Wide Association Analyses of Individual Cancers

We used PLINK^74, 75^ to implement within-cohort logistic regression models of additively modeled SNPs genome-wide, comparing cases of each cancer type to cancer-free controls. All models were adjusted for age at specimen collection, sex (non-sex-specific cancers only), first ten ancestry PCs, genotyping array (UKB only), and reagent kit used for genotyping (Axiom v1 or v2; GERA only). Case counts ranged from 471 (pancreatic cancer) to 13,903 (breast cancer) in the UKB and from 162 (esophageal/stomach cancer) to 3,978 (breast cancer) in GERA (**Supplementary Table 11**). Control counts were 359,825 (189,855 female) and 50,525 (29,801 female) in the UKB and GERA, respectively. After separate GWAS were conducted in each cohort, association results for the 7,846,216 SNPs in both cohorts were combined via meta-analysis. For variants that were only examined in one cohort (22% of the total 10,003,934 SNPs analyzed), original summary statistics were merged with the meta-analyzed SNPs to create a union set of SNP statistics for each cancer for use in downstream analyses (**Supplementary Figure 5**).

To determine independent signals in our union set of SNPs, we implemented the LD clumping procedure in PLINK^74, 75^ based on genotype hard calls from a reference panel comprised of a downsampled subset of 10,000 random UKB participants. For each cancer separately, LD clumps were formed around index SNPs with the smallest *P* not already assigned to another clump. To identify all potential signals, in each clump, index SNPs had a suggestive association based on *P*<1×10^−6^, and SNPs were added if they were marginally significant with *P*<0.05, were within 500kb of the index SNP, and had *r^2^*>0.1 with the index SNP. To confirm independence, we implemented GCTA’s conditional and joint analysis (COJO) method with the aforementioned downsampled subset of UKB participants as a reference panel,^76, 77^ performing stepwise selection of the index SNPs within a +/- 1000kb region of one another. SNPs were deemed independent if they maintained *P*<1×10^−6^ in the joint model. The remaining independent variants were determined to be novel if they were independent of previously reported risk variants in European ancestry populations (as described below).

To identify SNPs previously associated with each cancer type, we abstracted all genome-wide significant SNPs from relevant GWAS published through June 2018. We determined that a SNP was potentially novel if it had LD *r^2^* < 0.1 with all previously reported SNPs for the relevant cancer based on both the UKB reference panel and the 1000 Genomes EUR superpopulation via LDlink.^78^ As an additional filter for novelty, we again used COJO^76, 77^ to condition each potentially novel SNP on previously reported SNPs for the relevant cancer using the UKB reference panel, and SNPs were not considered novel if they did not maintain *P*<1×10^−6^ in the joint model. To confirm novelty and consider pleiotropy, we conducted an additional literature review to investigate whether these SNPs had previously been reported for the same or other cancers, including those not attaining genome-wide significance and those in non-GWAS analyses. For this additional review, we used the PhenoScanner database^79^ to search for SNPs of interest and variants in LD in order to comprehensively scan previously reported associations. We then supplemented with more in-depth PubMed searches to determine if the genes in which novel SNPs were located had previously been reported for the same or other cancers. Finally, for cancers with publicly available summary statistics (breast [>120,000 cases],^38^ prostate [∼80,000 cases],^80^ and ovarian [∼30,000 cases]^39^), we tested our potentially novel SNPs with *P*<1×10^−6^ for replication (defined as having the same direction of effect and *P*<0.05). Tested SNPs that did not replicate were not considered novel.

We considered whether clinical characteristics of the cases were informative about associated phenotypes by examining SEER stage and grade (GERA only) and age at cancer diagnosis (UKB and GERA). For each clinical variable, we decomposed cases into one of two categories: grade 1-2 (well or moderately differentiated) or grade 3-4 (poorly or undifferentiated); stage 0-1 (in situ or localized) or stage 2-7 (regional or distant metastases); age < median or age ≥ median. The case counts for all cancer-outcome strata are tabulated in **Supplementary Table 12**. For each of the novel GWAS SNPs, we conducted logistic regression comparing controls to each of the relevant case subtypes. We then compared the effect estimates across the strata for each clinical variable (e.g., for each relevant SNP-cancer pair, we compared the OR for grade 1-2 with the OR for grade 3-4) and calculated Cochran’s Q statistic to test for heterogeneity, adjusting for multiple testing for the number of strata and SNPs tested.

To assess whether our results were influenced by factors associated with survival, we conducted sensitivity analyses restricted to incident cases in the larger UKB cohort. For each cancer, we compared the independent SNPs that were suggestively associated in the analysis using both prevalent and incident cases (*P*<1×10^−6^) with those in the incident only analysis. We assessed whether the effect sizes varied by calculating Cochran’s Q statistic to test for heterogeneity, adjusting for multiple testing across the number of SNPs tested for each cancer. Additional sensitivity analyses evaluated esophageal and stomach cancers as separate phenotypes in the UKB cohort. For independent SNPs with *P*<1×10^−6^ in the analysis of the composite phenotype in UKB alone, we compared effect sizes for the composite phenotype to effect sizes for esophageal and stomach cancers separately and calculated Cochran’s Q statistic to test for heterogeneity, adjusting for multiple testing across the number of SNPs tested.

### Genome-Wide Heritability and Genetic Correlation

We used LD score regression (LDSC) on summary statistics from the union set of all SNPs genome-wide to calculate the genome-wide liability-scale heritability of each cancer type and the genetic correlation between each pair of cancer types.^81, 82^ Internal LD scores were calculated using the aforementioned downsampled subset of UKB participants. To convert to liability-scale heritability, we adjusted for lifetime risks of each cancer based on SEER 2012-2014 estimates (**Supplementary Table 13**).^83^ LDSC was unable to estimate genetic correlations for testicular cancer with both oral cavity/pharyngeal and pancreatic cancers, likely due to small sample sizes.

### Locus-Specific Pleiotropy

Using our union set of SNP-based summary statistics, we constructed pleiotropic regions of SNPs associated with more than one cancer with *P*<1×10^−6^. Non-overlapping regions were iteratively formed around index SNPs associated with any cancer, beginning with the SNP associated with the smallest *P*. We used a suggestive threshold of *P*<1×10^−6^ to assess whether suggestive regions for one cancer might also be informative for another. SNPs were added to a region if they were associated with any cancer with *P*<1×10^−6^, were within 500kb of the index SNP, and had LD *r^2^*>0.5 with the index SNP. We used a larger threshold for assessing pleiotropic regions (*r^2^*>0.5) than for identifying truly independent signals (*r^2^*>0.1; above) to ensure that all SNPs within a region were in LD. If all SNPs in a region were associated with the same cancer, the region was not considered pleiotropic.

### Genome-wide Variant-Specific Pleiotropy

We quantified one-directional and, separately, bidirectional variant-specific pleiotropy via the R package ASSET (*as*sociation analysis based on sub*sets*).^84^ Briefly, ASSET explores all possible subsets of traits for the presence of association signals, resulting in the best combination of traits to maximize the test statistic.^84^ ASSET has two procedures: in one, all traits are assumed to be associated with a variant in the same effect direction (one-directional pleiotropy); in the other, variants can be associated with traits in opposite directions (bidirectional pleiotropy).^84^ In the one-directional pleiotropy analysis, an overall *P* across the selected traits is provided, and in the bidirectional pleiotropy analysis, a *P* for each direction is provided as well as an overall *P* for the total association signal for both directions combined. ASSET corrects for the internal multiple testing burden accrued by iterating through all possible trait subsets for each variant as well as controlling for shared samples among the traits.^84^

Genome-wide ASSET analyses were conducted on the union sets of summary statistics for all 18 cancers. Independent variants were determined via LD clumping, where index SNPs were suggestively significant (overall *P*<1×10^−6^), and other SNPs were clumped with the lead variant if they had overall *P*<0.05, were within 500kb of the index SNP, and had *r^2^*>0.1 with the index SNP. We used a suggestive significance threshold to comprehensively assess all potentially pleiotropic variants. A SNP was determined to have a one-directional pleiotropic association if the overall *P* was <1×10^−6^ and it was associated with at least two cancers. A SNP was determined to have a bidirectional pleiotropic association if the overall *P* was <1×10^−6^ and the *P* for each direction was <0.05. For one- and bidirectional SNPs in LD with each other, the SNP with the smaller overall *P* was retained. We deconstructed bidirectional associations into cancers with risk-increasing effects and cancers with risk-decreasing effects.

To assess whether clinical aspects of the cases could be informative about the pleiotropic variants, for each of the one-directional and bidirectional pleiotropic SNPs, we conducted logistic regression comparing controls to each of the relevant case subtypes described above and calculated Cochran’s Q statistic to test for heterogeneity between estimates across the strata for each clinical variable.

### Functional Characterization of Pleiotropic Variants

Functional consequences for the 158 pleiotropic variants identified in the ASSET analysis were obtained from ANNOVAR. Enrichment of functional classes was evaluated using Fisher’s exact test, comparing the distribution observed among the pleiotropic variants to that of all variants in the reference panel of UKB European descent individuals.

Overall functional significance was assessed using DeepSEA,^33^ a deep learning tool that prioritizes functional variants by integrating regulatory binding and ENCODE modification patterns of ∼ 900 cell-factor combinations with evolutionary conservation features. Resulting functional significance scores, ranging from 0 to 1, represent the degree of deviation from a reference distribution of 1000 Genomes variants, with lower scores indicating a higher likelihood of functional significance. We also report CADD scores, which combine over 60 diverse annotations to predict deleteriousness.^35^ CADD scores are transformed into a log10-derived rank score based on the genome-wide distribution of scores for 8.6 billion single nucleotide variants in GRCh37/hg19 (i.e.: CADD=10 corresponds to top 10% most deleterious substitutions).^35^

To assess more specific functional features, we annotated each SNP according to Roadmap’s 15-core chromatin states across 127 cell or tissue types.^34, 85^ Chromatin state was assigned by taking the most common state across 127 cell or tissue types, with values ≤7 indicating open, accessible chromatin regions. Three-dimensional chromatin interactions were explored to identify significant interaction and enhancer-promoter links. Lastly, we explored associations with gene expression in blood and non-neurological tissues (since we did not investigate brain tumors) using data from the GTEx v7^86^ and BIOS QTL^59^ databases.

We conducted pathway analyses based on gene set collections from the Molecular Signatures Database (MSigDB)^87^ with an FDR *q*<0.05 significance threshold. Pathway enrichment analyses focused on the C2 MSigDB collection, which includes curated signatures of genetic and chemical perturbations, as well as canonical pathways from BioCarta, KEGG, Reactome, and other databases.

### Ethics

The study was approved by the University of California and KPNC Institutional Review Boards and the UKB data access committee, and informed consent was obtained from all participants.

## Data Availability

Our GWAS summary statistics are publicly available via direct request and will be deposited into dbGaP. The UKB cohort data is publicly available from the UKB access portal at https://www.ukbiobank.ac.uk. The UKB cancer phenotyping we performed to define cases will be provided to the UKB for public use. The Kaiser Permanente data are available via application with a local collaborator at: https://researchbank.kaiserpermanente.org/our-research/for-researchers/.

## Acknowledgements

This research was supported by the following National Institutes of Health grants: R01CA088164, R01CA201358, R25CA112355, K07CA188142, K24CA169004, and U01CA127298, and the UCSF Goldberg-Benioff Program in Cancer Translational Biology. The UK Biobank analyses were conducted using the UKB Resource under application number 14105. Support for participant enrollment, survey completion, and biospecimen collection for the RPGEH was provided by the Robert Wood Johnson Foundation, the Wayne and Gladys Valley Foundation, the Ellison Medical Foundation, and Kaiser Permanente national and regional community benefit programs. Genotyping of the GERA cohort was funded by a grant from the National Institute on Aging, the National Institute of Mental Health, and the NIH Common Fund (RC2 AG036607).

We thank the Breast Cancer Association Consortium (BCAC) for breast cancer summary statistics (http://bcac.ccge.medschl.cam.ac.uk/bcacdata/oncoarray/gwas-icogs-and-oncoarray-summary-results/), the Ovarian Cancer Association Consortium (OCAC) for ovarian cancer summary statistics (http://ocac.ccge.medschl.cam.ac.uk/data-projects/results-lookup-by-region/), and the Prostate Cancer Association Group to Investigate Cancer Associated Alterations in the Genome (PRACTICAL) consortium for prostate cancer summary statistics (http://practical.icr.ac.uk/blog/?page_id=8164).

## Author Contributions

S.R.R. and R.E.G. contributed by designing presented idea, conducting the analyses, and writing the manuscript. L.K. contributed to writing the manuscript. K.K.T. contributed by conducting analyses. S.E.A., M.A.B., T.B.C., D.A.C., N.C.E., J.D.H., E.J., L.H.K., T.J.M., S.K.V., E.Z., L.A.H., and T.J.H. aided in data acquisition, provided critical feedback, and helped shape the research, analysis, and manuscript. L.C.S. and J.S.W. contributed to study conception and design, supervised the project, and writing the manuscript.

