## Supplementary Figures 1-5 for "Pan-Cancer Study Detects Novel Genetic Risk Variants and Shared Genetic Basis in Two Large Cohorts"

**Supplementary Figure 1.** Comparison of effect sizes (log odds ratios) for associations between variants and cancers from incident case only analysis versus those obtained using both incident and prevalent cases. For all cancers, we compared associations for independent SNPs with $P<1x10^{-6}$ in the analysis with incident and prevalent cancers in UKB (2-95 SNPs per cancer). The effect estimates did not exhibit heterogeneity (P > 0.05/[number of SNPs per cancer]) and were highly correlated ($r^2$=0.95 across all 396 SNPs tested).

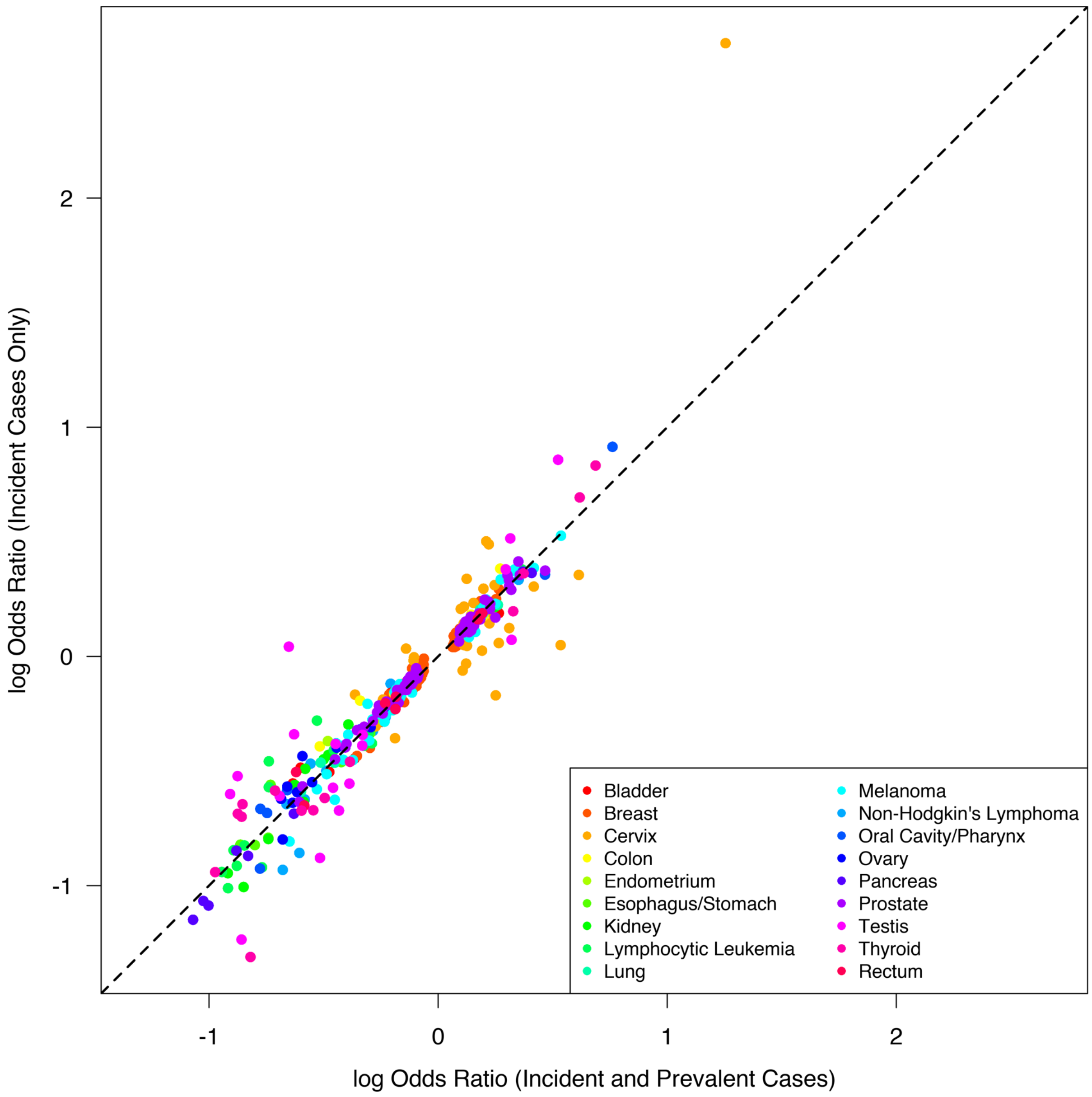

**Supplementary Figure 2**. Comparison of effect sizes (log odds ratios) for associations between variants and esophageal (left) and stomach (right) cancers versus those obtained using both cancers. We compared associations for independent SNPs with $P<1x10^{-6}$ in the analysis of the combined cancer phenotype in UKB alone (6 SNPs). The effect estimates did not exhibit heterogeneity (P > 0.05/6) and were highly correlated ($r^2$=0.98 comparing esophageal to combined and $r^2$=0.83 comparing stomach to combined).

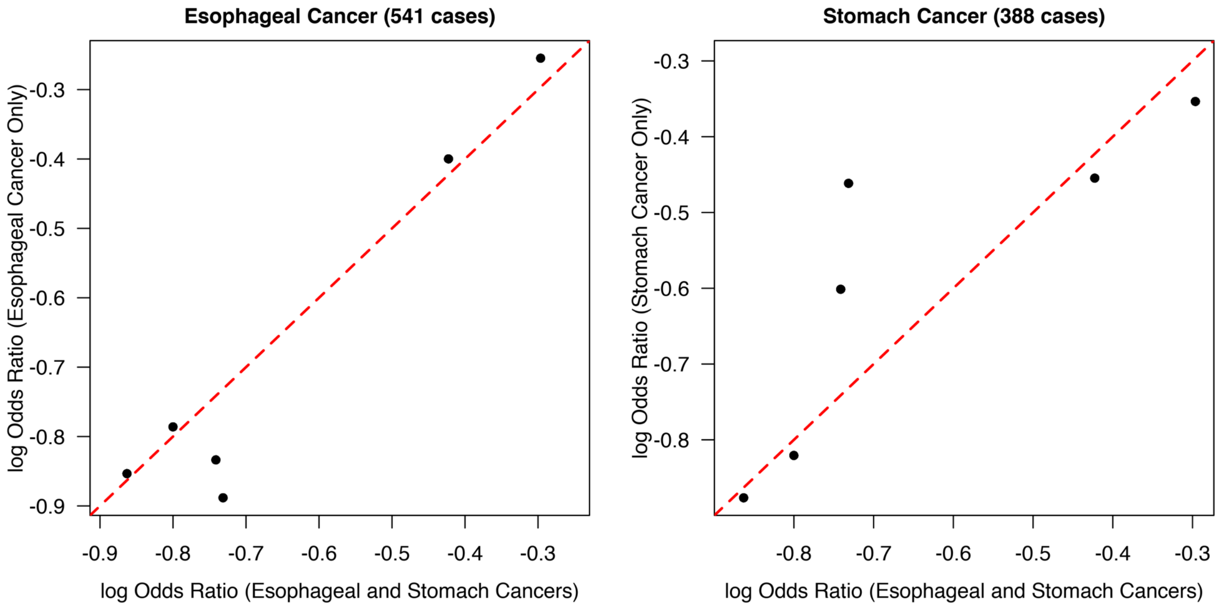

**Supplementary Figure 3.** Overview of significant (FDR<0.05) effects on gene expression observed for pleiotropic variants in BIOS-QTL and GTEx data sets. Histograms showing distribution of eQTL effects for (a) tissues and (b) genes across 83 pleiotropic variants with significant effects on gene expression. Distribution of tissues for 586 variant-gene pairs (c).

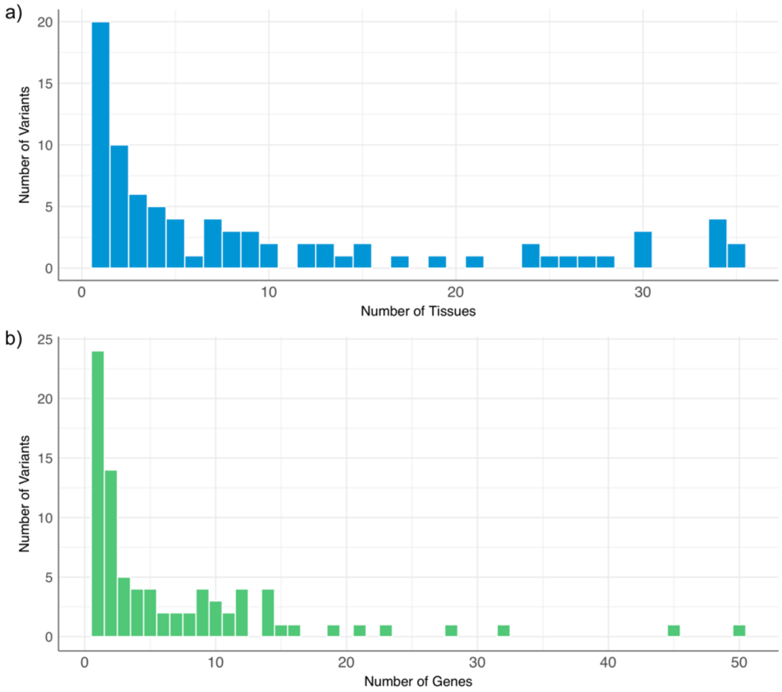


c)

| Tissue | N Genes | (%) |
|---|---|---|
| Whole Blood (BIOS QTL) | 359 | 61.3% |
| Adipose - Subcutaneous | 58 | 9.9% |
| Esophagus (Mucosa, Muscularis) | 31 | 5.3% |
| Muscle Skeletal | 17 | 2.9% |
| Artery (Aorta, Tibial) | 16 | 2.7% |
| Thyroid | 14 | 2.4% |
| Whole Blood (GTEx) | 11 | 1.9% |
| Cells Transformed fibroblasts | 9 | 1.5% |
| Lung | 9 | 1.5% |
| Heart | 8 | 1.4% |
| Skin | 8 | 1.4% |
| Adrenal Gland | 7 | 1.2% |
| Testis | 7 | 1.2% |
| Nerve - Tibial | 6 | 1.0% |
| Colon | 6 | 1.0% |
| Pancreas | 5 | 0.9% |
| EBV-transformed lymphocytes | 4 | 0.7% |
| Breast Mammary Tissue | 2 | 0.3% |
| Small Intestine Terminal Ileum | 2 | 0.3% |
| Spleen | 2 | 0.3% |
| Stomach | 2 | 0.3% |
| Gastroesophageal Junction | 1 | 0.2% |
| Prostate | 1 | 0.2% |
| Vagina | 1 | 0.2% |

**Supplementary Figure 4.** Circos plots depicting three-dimensional chromatin interactions (orange bands), such as HiC loops and enhancer-promoter links, observed for pleiotropic variants on chromosomes 11 and 22. All variants and links are plotted, but only variants in blue (bold) have significant effects.

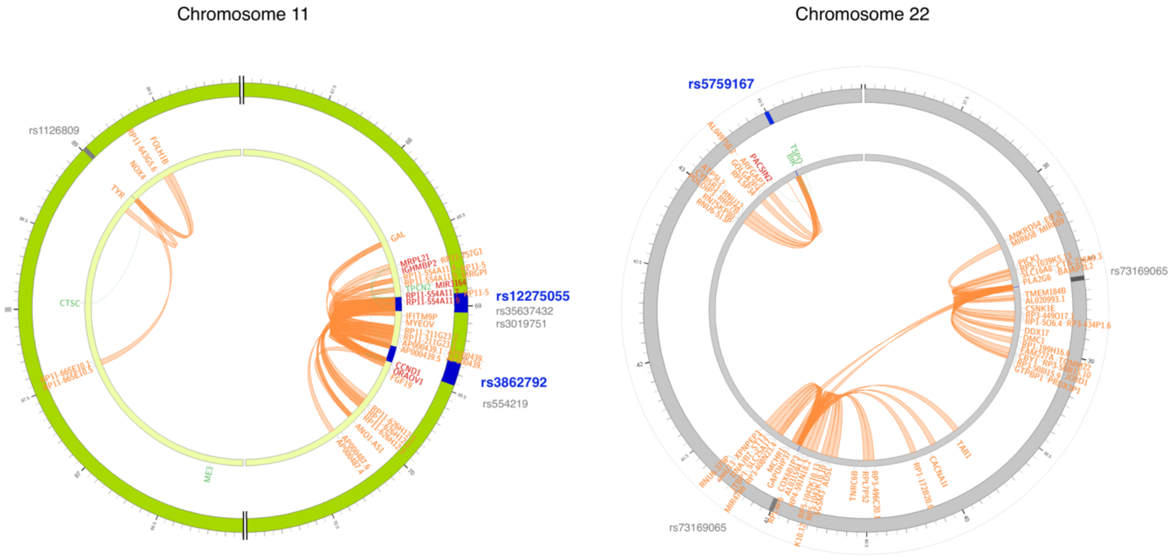

**Supplementary Figure 5.** Flowchart of overall approach used here. First we undertook individual cancer genome-wide association studies (GWAS) in two cohorts (UK Biobank [UKB] and Kaiser Permanente Genetic Epidemiology Research on Adult Health and Aging [GERA]). Based on those results, we used four approaches to assess: 1) novel risk variants; 2) heritability and genetic correlation; 3) locus-specific pleiotropy; and 4) variant-specific pleiotropy.

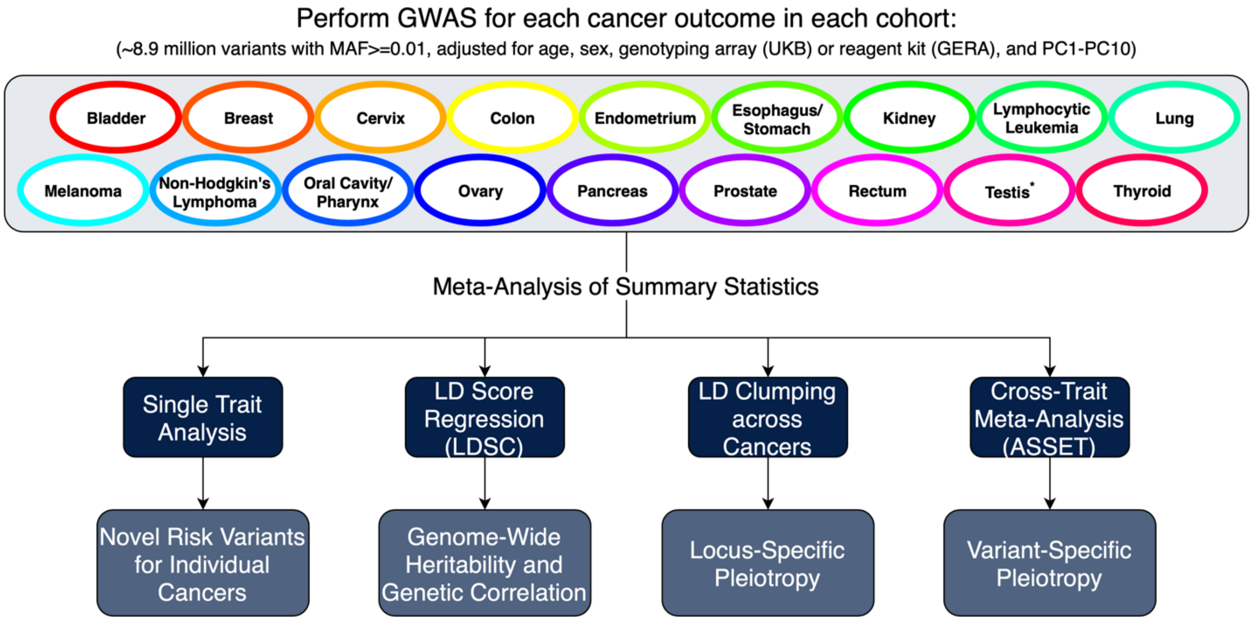


*Testicular cancer only analyzed in UKB
